# Rapid Adaptation to Road Salts in a Freshwater Microbial Eukaryote

**DOI:** 10.1101/2025.09.11.675700

**Authors:** Rebecca A. Zufall, Nia Pereda, Karissa Plum, Ethan Rothschild

## Abstract

Humans are changing habitat for wildlife in myriad ways and for populations to persist, they must adapt to this change. In parts of the world that experience snow and ice, road salts are often used to make driving safer in the winter. Runoff from these roads increases the salinity in nearby bodies of water, which has been shown to have detrimental physiological and ecological effects in freshwater ecosystems; however, the evolutionary consequences of salinization remain unclear. *Tetrahymena* are microbial eukaryotes that live in freshwater habitats and serve as an important link in the microbial food loop. In this study, we test how *T. thermophila* can evolve in response to increasing concentrations of road salts in their environment. Using experimental evolution, we found that *T. thermophila* adapt quickly to survive and grow better in increasing salinity. However, populations adapted to the highest salt concentrations experience fitness tradeoffs in salt-free environments. These results demonstrate the rapidity with which microbial populations can respond to anthropogenic changes to their environment, yet highlight the potential costs associated with this adaptation.

## Introduction

Human activity causes a variety of types of disturbance resulting in novel environmental conditions that pose challenges to the organisms in that environment. If and how populations will adapt to such rapid environmental change is an open question with important implications for biodiversity and ecosystem services (Peters et al., 2013; Reid et al., 2019). One important source of human-caused environmental disturbance comes from the use of road deicing salts in regions of the world that experience cold winters. Application of road salts improves road safety but has led to dramatically increased salinization of freshwater ecosystems (Dugan and Arnott, 2023; Hintz et al., 2022). In North America, the use of road salts started in the 1940’s and has been increasing each year since (Corsi et al., 2010; Kaushal et al., 2005). The effects of increased salinity in freshwater ecosystems include acute and chronic toxicity, changes in population dynamics, and reduced biodiversity (Corsi et al., 2010; Kaushal et al., 2005; Searle et al., 2016; Szklarek et al., 2022).

Several studies have examined the immediate impact of road salts on individuals, populations, and communities, demonstrating the potential for natural selection on salt tolerance (e.g. Hopkins et al., 2013; Schuler et al., 2017; Searle et al., 2016). Fewer studies have directly tested whether freshwater populations will adapt to increased salinity. Coldsnow et al. (2017) and Hintz et al. (2018) demonstrated that *Daphnia* populations rapidly evolve increased tolerance to road salts. Evidence of local adaptation in road-adjacent populations of salamanders (Brady, 2012) is also consistent with evolved tolerance to road salts. In contrast, Huber et al. (2024) found no increase in fitness in *Daphnia pulex* following evolution in salt. To further elucidate the long-term consequences of salinization of freshwater ecosystems due to road salts, it is necessary to better understand the potential for populations of various organisms to evolve in these novel conditions.

Ciliates are a diverse group of microbial eukaryotes that inhabit nearly every ecosystem worldwide (Lynn, 2012). In addition to their abundance, their unusual genome structure and diverse life history strategies have made them the subject of many studies. Ciliates are a critical link in the microbial food loop, facilitating energy flow from bacteria to larger organisms (Esteban and Fenchel, 2021; Weisse, 2017). Thus, to understand how freshwater ecosystems broadly will respond to salinization, it is important to determine how ciliates will respond. *Tetrahymena* is a well-studied genus of ciliates that are found worldwide in freshwater habitats (Doerder, 2019). *Tetrahymena* are primary predators that feed largely on bacteria and are preyed upon by zooplankton and small vertebrates (Lynn, 2008). *T. thermophila* is endemic to ponds and streams in the northeastern United States (Zufall et al., 2013), where road salt is frequently applied in the winter, making it a valuable system in which to study adaptation to increasing salinity.

Here, we tested how a freshwater microbial species would evolve in response to increasing salinity to further assess the ecological and evolutionary consequences of road salt application to freshwater ecosystems. We use experimental evolution to assess the effects of road salts on populations of *T. thermophila* (Plum et al., 2022). We ask whether and how rapidly populations can adapt to tolerate increasing salt concentrations, whether there are tradeoffs for salt-adapted populations in salt-free environments, and whether adaptation to one salt increases fitness in other types of salt.

## Materials and Methods

### Study System and culture conditions

*Tetrahymena thermophila* strain SB210E (*Tetrahymena* Stock Center ID SD01539) was used for all experiments. Cells were grown in a standard nutrient-rich *Tetrahymena* medium, SSP, with added penicillin-streptomycin-amphotericin solution (Cassidy-Hanley, 2012; Gorovsky et al., 1975) for all experiments, either with or without added salts as described below. 100% SSP was used for growth conditions and 5% SSP was used for population maintenance with minimal cell division.

Prior to the start of the experiment, *T. thermophila* was assayed for salt tolerance. 200 µl of cells were put into replicate wells with 0, 3, 6, 9, 12, or 15 g/L of NaCl or MgCl_2_ in SSP. At 0.25, 0.5, 1, 2, 24, and 48 hours following salt exposure, cells were observed under a microscope for movement and cell division. Dose response curves were plotted as the salt concentration vs. fraction of cells moving, and the IC50 was estimated as the concentration at which 50% of cells were not moving after 24 hours. These IC50 values were used as the starting concentrations of salt during experimental evolution. We later discovered that cell movement is not a good proxy for survival, thus these IC50 values do not correspond to the IC50 values we measured at the end of the experiment. Nonetheless, they served as useful starting conditions for the evolution experiment.

### Experimental Evolution

Prior to the start of the experiment, *T. thermophila* were thawed from liquid nitrogen storage and a single cell was isolated and grown to stationary phase for one week in SSP at room temperature. 300 µl of cells were transferred to wells of a 12-well plate, in 3.5 mL of SSP, containing either no added salt (this was the control for adaption to the laboratory conditions), 9 g/L NaCl, or 5 g/L MgCl_2_. The salt treatments were either “constant,” i.e. the same concentration of salt was used throughout the experiment (designated N and M for constant NaCl or MgCl_2_, respectively), or “increasing,” i.e. the salt concentration was increased each week of the experiment until the populations could no longer adapt (designated N+ and M+; Table 1). The N+ lines reached a concentration of 21 g/L NaCl after 62 days, but then all populations died. These populations were thus restarted from backup populations that had been adapting to 9 g/L NaCl for 85 days, then took an additional 17 days to reach 18 g/L NaCl. The NaCl concentration was not increased beyond 18 g/L so cell lines would not be lost again. (Note that the original N+ populations reached 18 g/L NaCl after 50 days.) Thus for NaCl, the final concentration in the increasing populations was 18 g/L. For MgCl_2_, the final concentration was 17 g/L, reached after 91 days. Each evolution treatment had four replicate populations. All cell culture was performed at 24°C in an incubator.

**Table 1.**
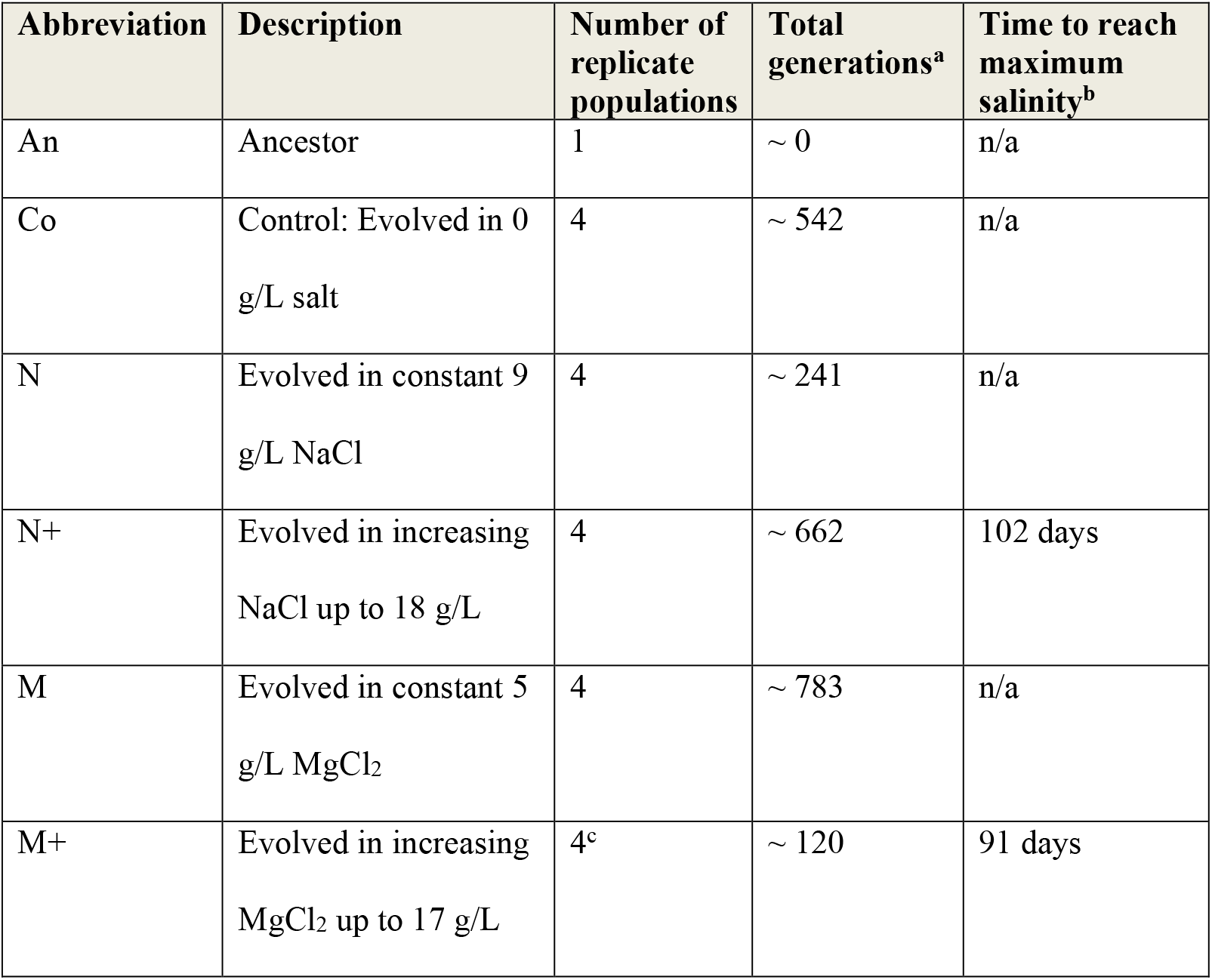
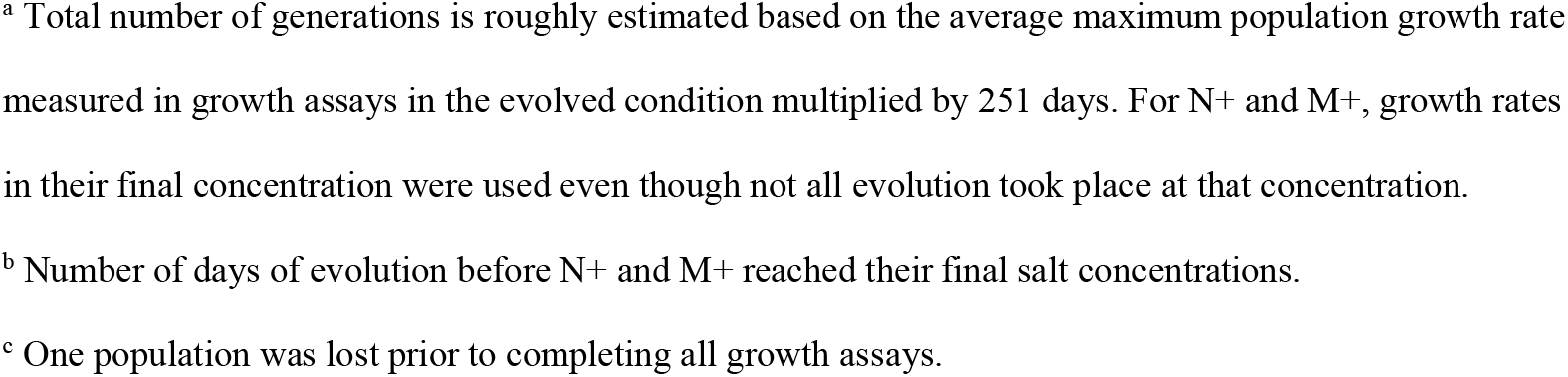
Evolution conditions for each experimental treatment.

| Abbreviation | Description | Number of replicate populations | Total generations<sup>a</sup> | Time to reach maximum salinity<sup>b</sup> |
| --- | --- | --- | --- | --- |
| An | Ancestor | 1 | ~ 0 | n/a |
| Co | Control: Evolved in 0 g/L salt | 4 | ~ 542 | n/a |
| N | Evolved in constant 9 g/L NaCl | 4 | ~ 241 | n/a |
| N+ | Evolved in increasing NaCl up to 18 g/L | 4 | ~ 662 | 102 days |
| M | Evolved in constant 5 g/L MgCl<sub>2</sub> | 4 | ~ 783 | n/a |
| M+ | Evolved in increasing MgCl<sub>2</sub> up to 17 g/L | 4<sup>c</sup> | ~ 120 | 91 days |

<sup>a</sup> Total number of generations is roughly estimated based on the average maximum population growth rate measured in growth assays in the evolved condition multiplied by 251 days. For N<sup>+</sup> and M<sup>+</sup>, growth rates in their final concentration were used even though not all evolution took place at that concentration.
<sup>b</sup> Number of days of evolution before N<sup>+</sup> and M<sup>+</sup> reached their final salt concentrations.
<sup>c</sup> One population was lost prior to completing all growth assays.

Each population was subcultured to a new well every 48-72 hours. Cells were thoroughly mixed via pipetting, then 10% was transferred to fresh medium for a final volume of 3.5 mL. At each transfer, cultures were visualized under the microscope to ensure ongoing cell division and no contamination. When contaminants (usually fungal) were observed, new subcultures were started from the most recent prior culture of that replicate that did not have a contaminant.

Populations were maintained in their salt treatments for a total of 251 days, or approximately 500 generations on average. Cell counts were not made at each subculture, so we estimated the elapsed number of generations based on the average maximum population growth rate of each population in its evolved condition (Table 1).

The ancestor was maintained in 5% SSP, which dramatically slowed cell division, but did not stop it. Thus, the ancestor we assayed is not identical to the original population used to start the experimental populations. Once the evolution experiment was completed all populations were moved to 5% SSP (plus the same/final concentration of salt as their evolution condition) or liquid nitrogen storage until assays could be performed.

### Growth assays

Following evolution, all populations were assayed for growth in each salt condition. Prior to starting a growth assay, all cells were acclimated to SSP with no salt for at least 2 days. Following acclimation, cell density was determined by counting under a microscope. Approximately 150 cells were transferred to wells in a 96-well plate containing 170µl of medium. Five assay plates were used containing the salt concentrations used in each evolution condition, or the final concentration for the increasing treatments: no salt (SSP), 9 g/L NaCl (Na9), 18 g/L NaCl (Na18), 5 g/L MgCl_2_ (Mg5), and 17 g/L MgCl_2_ (Mg17). Cells from each of the evolved populations, and the ancestor, were randomized on each plate in 3 wells each (for triplicate measurement replication), except the ancestor, which had 12 replicates. The remaining wells contained media. Prior to completing the two MgCl_2_ assays, one of the M+ populations was lost. OD_650_ was measured on a microplate reader every 10 minutes, with shaking, for 6 days. Following completion of the assay, all wells were inspected microscopically to ensure no contamination.

### Data analysis

#### Growth curve fitting

Growth curves were analyzed using the R package growthrates (Petzoldt, 2022). Because we observed that lag time varies greatly among treatments, we only fit growth models that contained a parameter for lag time. The Huang (2011) model provided the best overall fit based on residual sum of squares. Three parameters were extracted from the fit curves: maximum growth rate, carrying capacity, and lag time.

#### Survival analysis

Growth models that fit the data with r^2^ > 0.90 were considered to have survived. Visual inspection of growth models confirmed that models with r^2^ < 0.90 did not fit well because OD values never increased above the detectable threshold, indicting little or no cell division in that well due to cell death. Cell death in these conditions was confirmed by staining with the vital dye trypan blue.

Survival probability was analyzed with a logistic model (survival ∼ evolution condition ^*^ assay condition) using brglm_fit. implementing mean bias reduction due to complete separation of the data (Kosmidis et al., 2020). Marginal means and contrasts were found using the R package modelbased (Makowski et al., 2025).

#### Dose response curves

For dose response curves, we combined data from the two salts and considered only chloride ion concentration (survival probability ∼ [Cl^-^]). The R package drc (Ritz et al., 2015) was used to fit dose response curves, extract IC50 values, and test for pairwise differences in IC50 values.

#### Growth parameter analysis

For assayed populations that survived, we further analyzed growth curve parameters. Populations that did not reach carrying capacity during the assay (visually identified as wells with carrying capacity > 0.8 or maximum growth rate > 0.25) were removed from the analysis. Linear mixed effect models were used to assess the effects of evolution condition and assay condition on the three growth parameters: carrying capacity, maximum growth rate, and lag time (growth parameter ∼evolution condition ^*^ assay condition + replicate population nested within evolution condition). The R package emmeans was used to estimate marginal means and perform pairwise comparisons (Lenth, 2025).

## Results

Replicate populations of *T. thermophila* were evolved under either constant or increasing concentrations of two of the most commonly used road salts, NaCl and MgCl_2_. After ∼500 generations, changes in salt tolerance of all populations (Table 1) were measured via population growth curves, which were used to estimate survival probability and growth parameters.

### Survival of evolved populations

Survival was defined by the ability of cells to grow in each salt concentration. Evolution condition, assay condition, and the interaction between these all significantly affected the probability of survival (ANOVA, p < 0.01). Survival was high for all populations in the no-salt (SSP) and medium-salt conditions (Na9 and Mg5), with two exceptions (Fig. 1). The ancestor had decreased survival in 9 g/L NaCl relative to the other populations (marginal contrasts, p < 0.05). In addition, the M+ populations had a marginally significant decrease in survival in the no-salt condition relative to the other populations (p = 0.1).

**Figure 1.**
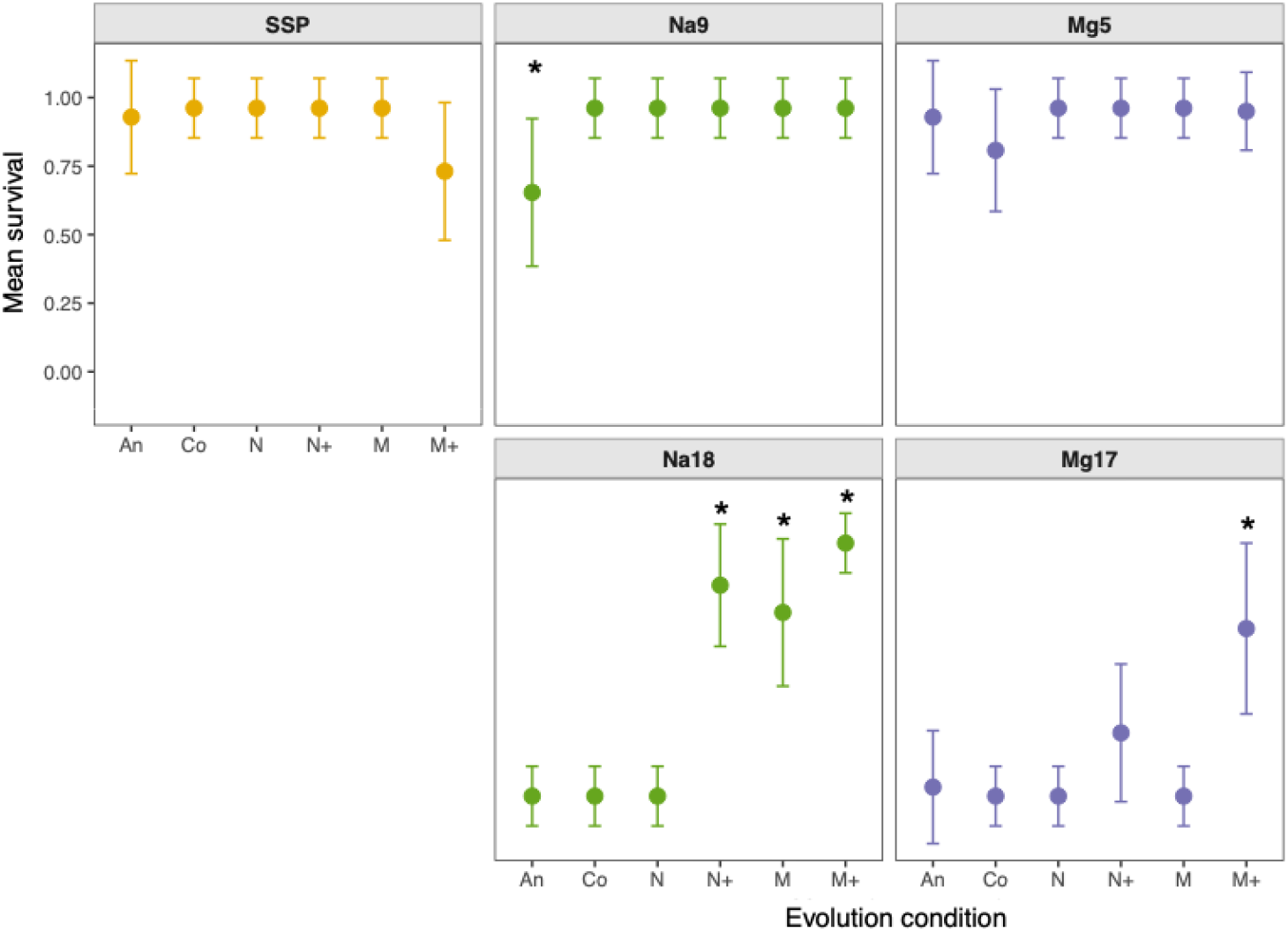
Survival of each population in no salt (SSP), medium salt, i.e. 9 g/L NaCl (Na9) or 5 g/L MgCl_2_ (Mg5), or high salt, i.e. 18 g/L NaCl (Na18) or 17 g/L MgCl_2_ (Mg17). Survival is the marginal mean of the replicate populations shown with 95% confidence intervals. Evolution conditions are as in Table 1. ^*^ indicates values that differ significantly from the others in marginal contrasts within assay condition (p < 0.05), except that M+ is not significantly different from N+ in Mg17.

In high-salt conditions (Na18 and Mg17), the increasing-salt evolved lines tend to survive much better than the others (Fig. 1). In 18 g/L NaCl, M, M+, and N+ all survive significantly better than the other populations (Fig. 1). In 17 g/L MgCl_2_, M+ survives better than all other lines, except N+, which also survives marginally better than the other lines (p = 0.1; Fig. 1).

### Evolved dose response curves

Combining the data from NaCl and MgCl_2_ assays, we determined half-maximal inhibitory concentrations (IC50) of chloride using dose response curves (Fig. 2). All evolved lines, including the control, had IC50 values trending higher than the ancestor. IC50 was significantly higher than the ancestor in N+ and M lines (t-test, p < 0.05). The estimate of IC50 is highest in the M+ line, however this estimate has a high degree of uncertainty because it had high survival rates even in the highest measured salt concentrations (Fig. 2).

**Figure 2.**
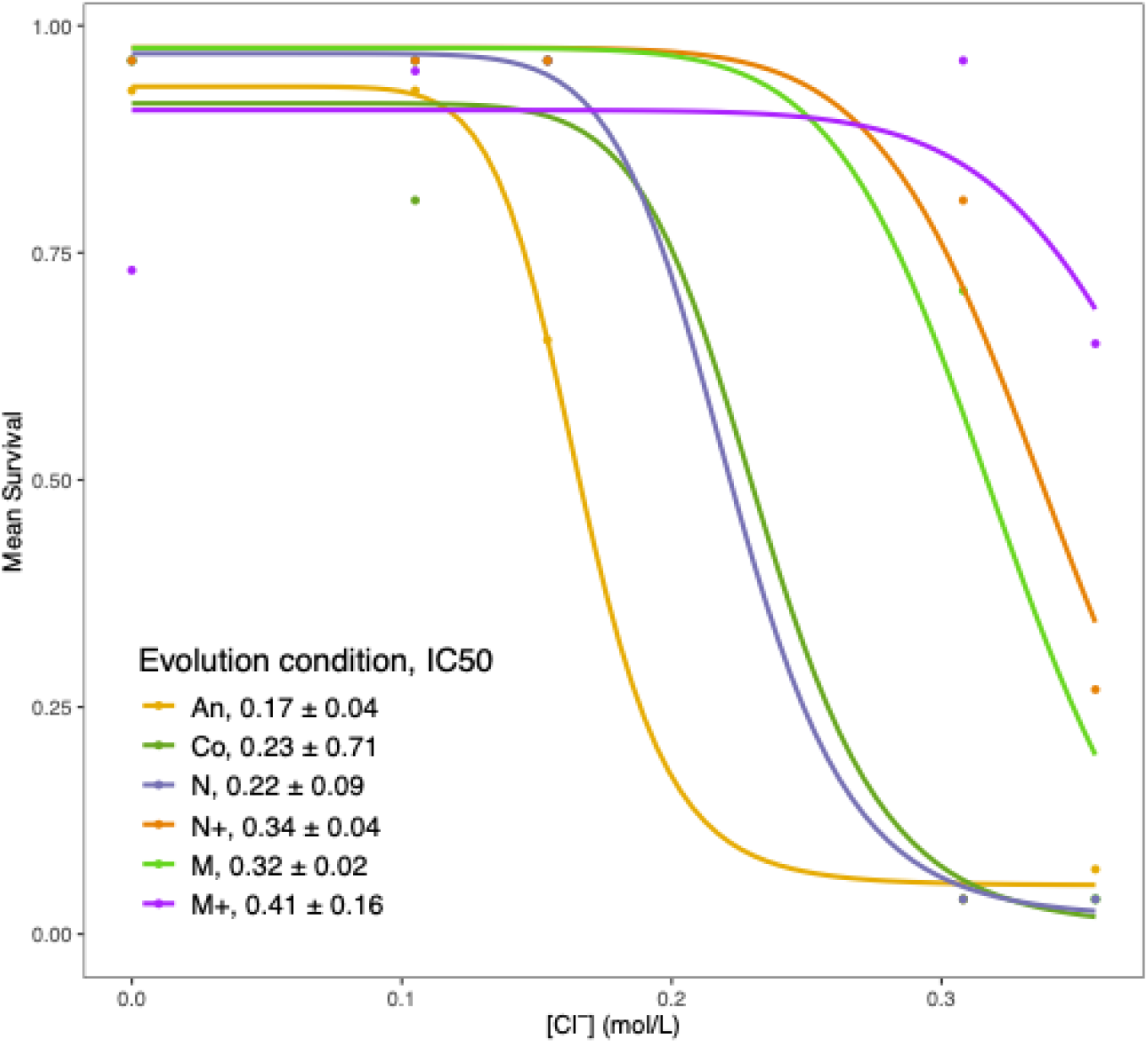
Dose response curves based on marginal means of survival probability in different chloride ion concentrations. Curves were fit using a four-parameter log-logistic function in the R package drc. IC50 values are shown +/-standard error.

### Effect of salt adaptation on growth parameters

Populations that survived were grown for 6 days. Growth curves were modeled and lag time, maximum growth rate, and carry capacity were estimated from these models. All three growth parameters were significantly affected by evolution condition, assay condition, and their interaction (ANOVA, p < 0.001).

The ancestor generally did not differ from most other lines when grown in 5 g/L MgCl_2_, but had significantly longer lag time and lower carrying capacity than all of the evolved lines in 9 g/L NaCl (marginal contrasts, p < 0.05; Fig. 3). This is consistent with fact that 5 g/L MgCl_2_ is below the IC50 of the ancestor and 9 g/L NaCl is approximately equal to the ancestor’s IC50.

**Figure 3.**
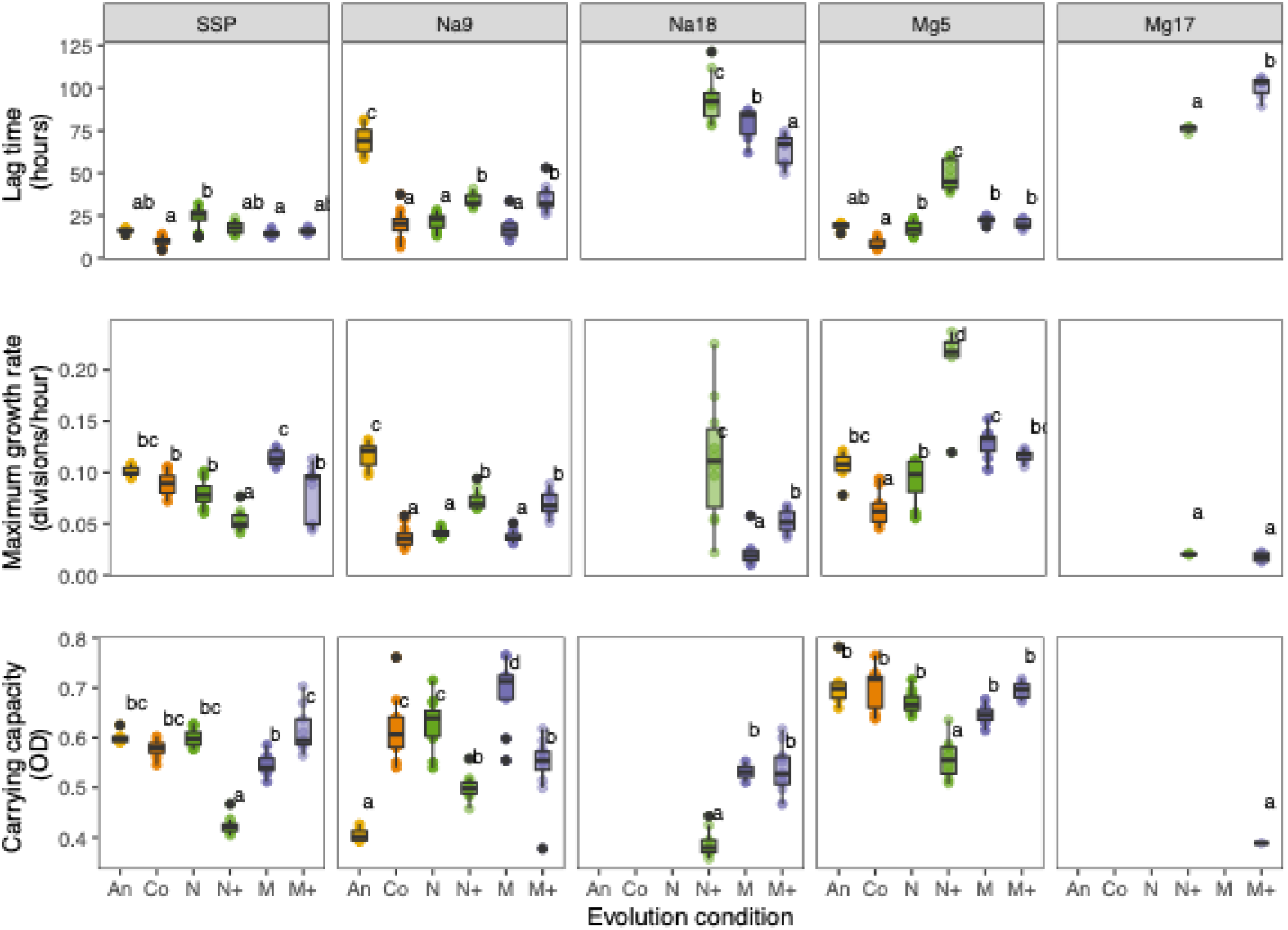
Growth parameters of surviving populations. Lag time, maximum population growth rate, and carrying capacity were estimated from growth curve models of each population assayed in each salt concentration. Abbreviations are as in Fig. 1. Pairwise comparisons of marginal means within in each assay condition are shown; values that share a letter are not significantly different from one another at p < 0.05.

Surprisingly, the maximum growth rate of the ancestor in 9 g/L NaCl is significantly higher than the other populations (marginal contrasts, p < 0.05; Fig. 3). The pattern of a long lag time and low carrying capacity in conjunction with a faster growth rate is also observed in the N+ populations when grown in the medium salt conditions and 18 g/L NaCl. Looking at all of the populations together, there is not a significant correlation between lag time and growth rate (Spearman rank correlation, R^2^ = -0.084, p = 0.21), but the ancestor and N+ populations both show a positive correlation between these parameters whereas all other populations show a negative relationship (Fig. 4).

**Figure 4.**
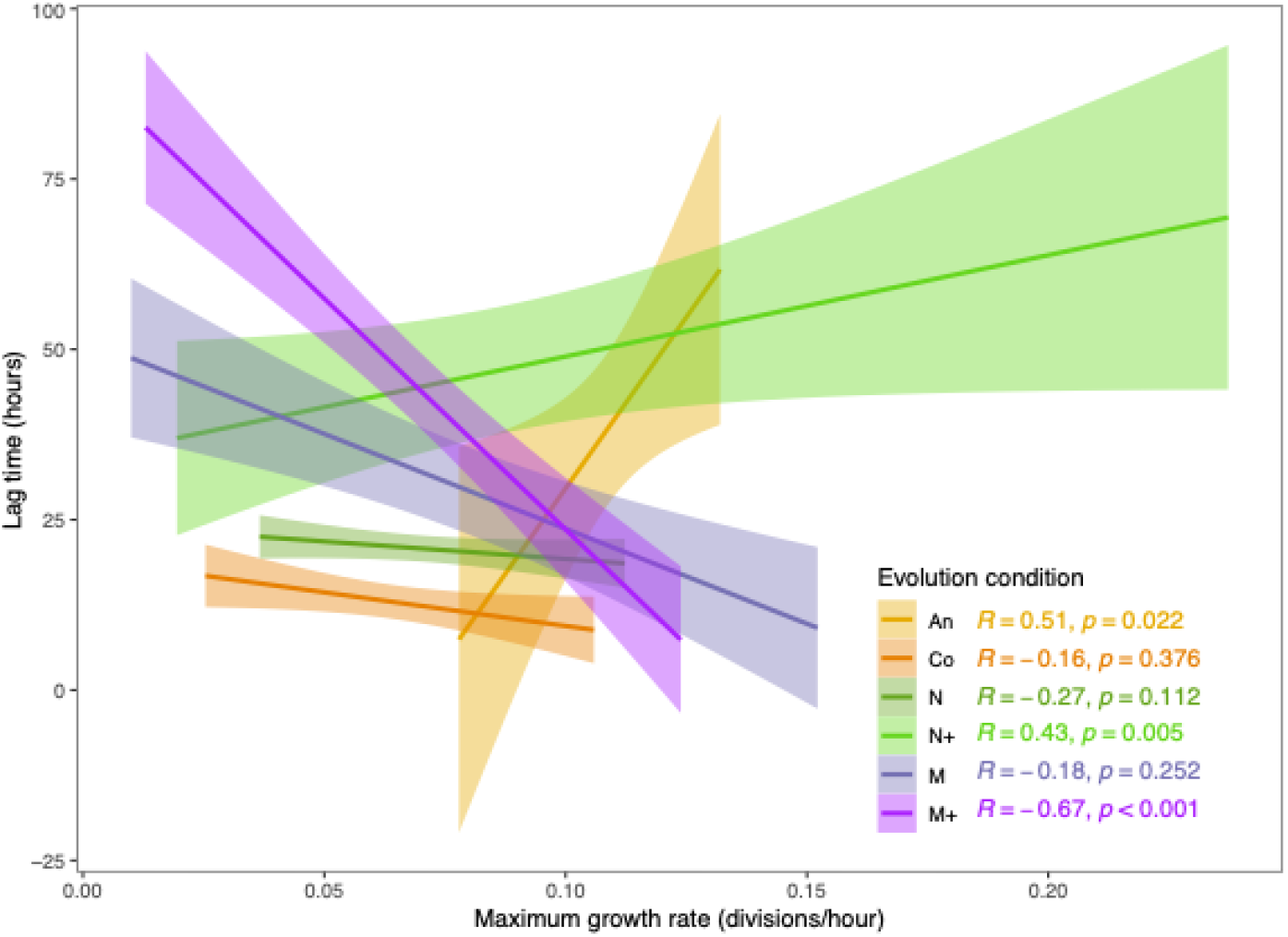
Correlation between lag time and growth rate differs between evolution treatments. Linear regression, 95% confidence interval, and Spearman correlation coefficient are shown for each evolution condition.

The control populations, i.e. those evolved in no added salts, outperformed the ancestor when grown in 9 g/L NaCl, with shorter lag times and higher carrying capacity (Fig. 3). They also survived slightly better and had higher IC50s than the ancestor (Fig. 1, 2). This suggests that adaptation to laboratory conditions increases salt tolerance even when not evolved in the presence of salt.

N+ and M+ populations evolved the highest salt tolerances, but this came with a cost when grown in no-salt or medium-salt concentrations. In particular, N+ populations had significantly reduced growth rate and carrying capacity compared to all other populations when grown in no salt (SSP), and longer lag times and lower carrying capacities than the control, N, and M evolved populations when grown in medium-salt concentrations (Fig. 3). M+ populations only show increased lag time and reduced carrying capacity in 9 g/L NaCl relative to the control, N, and M populations. In addition, slightly fewer M+ populations survived in SSP (Fig. 1), however those that survived exhibited growth patterns in SSP similar to the other populations (except N+; Fig. 3).

## Discussion

Freshwater ecosystems are experiencing detrimental effects of increased salinization due to application of road salts (Dugan and Arnott, 2023; Hintz et al., 2022). Determining how populations will adapt to this environmental challenge is critical for understanding the future health of these habitats. Here, we demonstrated that *T. thermophila* has the potential to adapt quickly to increasing concentrations of two of the most commonly used road salts. Following evolution in salt, ciliate populations were able to survive in significantly higher salt concentrations than the ancestral population. This result is consistent with previous studies in *Daphnia*, which were also shown to quickly evolve increased tolerance to road salts (Coldsnow et al., 2017).

Considering these results in the context of wild populations, measured concentrations of Cl^-^ in freshwater systems in the northeastern United States have been reported to reach as high as 0.03-0.13 mol/L (Brady and Benoit, 2025; Huber et al., 2024; Kaushal et al., 2005), close to the IC50 of the *T. thermophila* ancestral population. This suggests that natural populations of ciliates are likely to experience similar selective conditions as those imposed by our Na9 (0.154 mol/L) and Mg5 (0.105 mol/L) treatments. In addition, the timescale of our experiment is short enough that there would be sufficient time in a single season to see the responses we observed in lab. In particular, the N+ and M+ lines adapted to survive in high salinity after only 50 (for the original N+ lines that were lost) or about 100 generations (Table 1). While we do not know how rapidly cells are dividing in natural conditions, it is likely that they undergo more than 100 generations in a single year and may experience similar salinities to those in our experiment, suggesting that the responses observed here are likely to be relevant to wild populations.

One factor from natural populations that was not taken into consideration in this experiment is the fact that application of road salts is not constant across time. Because road salts are applied in winter months, the highest salinity levels tend to be found in winter and early spring, with salinity decreasing through summer and fall (Brady and Benoit, 2025; Dugan and Arnott, 2023; Kaushal et al., 2005). These fluctuations in salinity levels are of particular interest given the observed fitness trade-offs in the lines adapted to the highest salt concentrations. Compared to the freshwater adapted ancestor, N+ and M+ lines evolved the highest salt tolerance, but had reduced survival (M+), reduced growth rate (N+), and longer lag times (M+ and N+) when grown in salt-free conditions. This suggests that adaptation to high salinity in spring can be costly when salinity is lower later in the year. Further experiments testing the effects of these annual fluctuations in salinity levels will be important to determine the long-term effects of road salts on these populations.

Tradeoffs, whereby increased fitness in one environment leads to decreased fitness in another, such as those we find in the N+ and M+ lines, have been found in a wide variety of situations across taxonomic groups (e.g. Agudelo-Romero et al., 2008; Gompert and Messina, 2016). However, positive correlated responses, where selection in one environment leads to increased fitness in another, have also been found in a variety of contexts (e.g. Magalhães et al., 2009; Nidelet and Kaltz, 2007; Olazcuaga et al., 2021; Tarkington and Zufall, 2024). Here, we find positive correlated responses to selection whereby evolution in NaCl increases fitness in MgCl_2_ and vice versa. Given the similarity of these environments, perhaps this correlation is not surprising, although previous studies have found different responses to different road salts (Huber et al., 2024). More surprising is the fact that the control lines, evolved in no salt, also showed a positive correlated response when grown in salt, with an IC50 nearly the same as the lines evolved in 9 g/L NaCl. The mechanism for this response is unclear but corresponds to results from a previous study on experimentally evolved lines of *T. thermophila*, which showed frequent positive correlated responses to selection where evolution in one environment led to increased fitness in other environments (Tarkington and Zufall, 2024). Here, it appears that adaptation to lab conditions has the correlated effect of increasing salt tolerance.

Considering a different kind of tradeoff, Huber et al. (2024) found life history tradeoffs in *Daphnia* grown in salt. When grown in low salt, *Daphnia* had decreased brood size but increased lifetime reproductive output due to longer life spans. In the An and N+ populations here, we observe a similar life history tradeoff, where long lag times are followed by high growth rates. The usual expectation for the relationship between lag time and growth rate is a negative correlation—low fitness genotypes will have both long lag times and low growth rates. This has been found extensively (e.g. Baranyi and Roberts, 1994; Basan et al., 2020) and is what we found in all of the populations besides An and N+. However, in the An and N+ populations an inverse relationship indicates that these populations tend to take a longer time to acclimate to a new condition, but once they do, they are able to tolerate it better and grow faster. It is interesting that the two populations that show this pattern have the lowest and one of the highest salt tolerances. This result may be related to the finding of (Basan et al., 2020) that cells with a high growth rate tend to have longer lag times following an environmental shift, however, the mechanism remains unclear.

Ciliate reproductive biology adds an additional complication to understanding how populations will adapt to increasing salinity. Ciliates contain two types of nuclei in each cell: a germline micronucleus, which is quiescent during asexual reproduction, and a somatic macronucleus, from which all gene expression is driven during growth and reproduction (Merriam and Bruns, 1988; Prescott, 1994). Thus, in this experiment, where all reproduction was asexual, adaptation occurred solely due to mutations in the macronucleus. However, following sexual conjugation, the macronucleus gets degraded and a new macronucleus will develop from a zygotic nucleus (the product of syngamy between two haploid micronuclei; Orias et al., 2011). This means that any mutations that were selected in the macronucleus during asexual division will be lost following sex. It remains unknown when or how frequently *T. thermophila* undergo sexual conjugation in the wild (Doerder et al., 1995; Lynn and Doerder, 2012; Zufall, 2016), however, it is clear that this will change the dynamics of adaptation in these populations. Further experiments are necessary to understand the effects of sex and generation of a new macronucleus on adaptation to increasing salinity.

## Data availability

All data are available on Dryad: https://doi.org/10.5061/dryad.vhhmgqp6v

## Acknowledgements

This work was supported by NSF DEB 2342961.

